# Genetic continuity in the last seven Millennia in human hepatitis B viruses

**DOI:** 10.1101/404426

**Authors:** Xiaoyun Lei, Ye Zhang, Shi Huang

## Abstract

Hepatitis B virus (HBV) is a major human pathogen and yet the evolution history of HBV has largely remained uncertain. With a better theoretical understanding of genetic diversity, we here used a new method to examine the previously published ancient and present day HBV genomes. We identified an informative region in the HBV polymerase that is slow evolving and used it to study genetic distances among HBVs. Three ancient human HBV isolates from 4488-7074 years ago in Germany were identified as genotype G that is also presently common in the same country. We constructed a new phylogenetic tree of HBVs that placed genotype D as the most basal branch with an inferred age of ~20500 years, which is remarkably consistent with the worldwide distribution and a most parsimonious migration route of HBV genotypes today. These results help resolve the evolutionary history of HBV and provide a useful method for studying the phylogenetics of HBV and other viruses in general.

## Introduction

Hepatitis B virus (HBV) is a major cause of human hepatitis and related diseases (http://www.who.int/mediacentre/factsheets/fs204/en/). The origin and evolution of HBV has largely remained uncertain, like most viruses. HBV has a circular, partially double-stranded DNA genome of about 3.2kbp that encodes four overlapping open reading frames (P polymerase, pre-S/S envelope, pre-C/C core protein, and X). At least 8 genotypes (A–H) based on nucleotide sequence similarity are classified for human HBV and they have a heterogeneous global distribution (Castelhano, Araujo, Arenas 2017). The putative basal genotypes F and H are found exclusively in the Americas, thus inconsistent with the notion that HBV co-evolved with modern humans as part of the Recent Out of Africa hypothesis. Yet, HBVs in non-human primates (NHP), such as chimpanzees and gorillas, are phylogenetically related to human HBV isolates, seemingly supporting the idea of an Africa origin of the virus (Locarnini et al. 2013; Souza et al. 2014).

Recently, a number of ancient HBV genomes have been uncovered from human skeletons found in Europe and Asia that are between approximately 500-7000 years ago (Kahila Bar-Gal et al. 2012; Krause-Kyora et al. 2018; Muhlemann et al. 2018). While most of the relatively younger HBV genomes (<4488 years ago) were closest to present day human HBVs, all three oldest HBV samples found in Germany (between 4488-7074 years ago) were unexpectedly closest to chimpanzee or gorilla HBVs and hence considered extinct today. The finding challenges expectations as HBV today must have an ancient ancestor which must have infected a large population in the past to have a chance to survive to the present. As large populations have a greater probability of having some of its remains discovered today, the probability of discovering an ancestor of today’s human HBVs should be much greater than that of finding a now extinct ancient human HBV sample. Thus, the unusually high rate of discovering ancient human HBV samples that are now extinct (3 independent samples in 3 different archaeological sites) indicates potential flaws in the phylogenetic method employed, especially given that existing methods have yet to produce a consistent evolutionary history of the HBVs. Importantly, the theoretical framework underlying the existing methods, the neutral theory, has been widely known to be inadequate as an explanatory theory of the observed genetic diversity patterns (Kreitman 1996; Ohta, Gillespie 1996; Hahn 2008; Leffler et al. 2012; Hu et al. 2013; Kern, Hahn 2018). It is unfortunate that existing phylogenetic methods have relied heavily on the neutral theory being a valid interpretation of nature.

Different positions in a viral genome are known to have different mutation rates. The fast changing sites in influenza virus play adaptive roles in escaping host immune defense and undergo constant and quick turnovers (Shih et al. 2007). The antigenic sites in human influenza A virus mutate and turn over quickly (several times within a 30 year period), which is critical for the virus to escape host immune defense and hence for flu epidemics. In contrast, other sites stayed largely unchanged within the same period. The influenza results illustrate two general points with regard to evolutionary dynamics of a genome that have so far been overlooked. First, fast evolving or less conserved DNAs are also functional rather than neutral as they are essential for quick adaptive needs in response to fast changing environments. Second, fast evolving DNAs turn over quickly and can be shown to violate the infinite sites model. Hence, they cannot be used for phylogenetic inference involving evolutionary timescales. If one uses the fast changing sites in a flu virus to infer the phylogenetic relationship of the virus isolates responsible for different epidemics in a past period of say 10 years, one would have reached the erroneous conclusion that each epidemic was caused by a distinct type of flu virus with no genetic continuity among them rather than just minor variations of the same type.

For short term lineage divergence that has yet to reach saturation for the fast changing sites, both fast and slow changing sequences could be informative to phylogeny. However, for evolutionary timescale where divergence in fast changing sites have reached saturation, only the slow sites (the slow clock method) could be informative, as has been previously shown and explained by the maximum genetic diversity (MGD) hypothesis (Huang 2012; Hu et al. 2013; Huang 2016; Yuan, Huang 2017; Yuan et al. 2017). The MGD hypothesis has recently solved the longstanding puzzle of genetic diversity (Huang 2009; Huang 2016) and made it now possible for the first time to realistically infer phylogenetic relationships based on genetic diversity data. It has now been demonstrated that genetic diversities are mostly at saturation level (Yuan et al. 2012; Yuan et al. 2014; Zhu et al. 2015a; Zhu et al. 2015b; Gui, Lei, Huang 2017; He et al. 2017; Lei, Huang 2017; Lei et al. 2018; Teske et al. 2018), which therefore renders most of the past molecular results invalid since those results were based on mistreating saturated phases of genetic distance as linear phases. Only slow evolving nuclear sequences are still at linear phase and hence informative to phylogenetic relationships.

Here, we investigated the genetic relationships among the ancient HBVs and present day human HBVs using the slow clock method. We found that all three ancient HBV samples that were thought to group with NHP isolates in fact grouped with human HBVs. We also constructed a new phylogenetic tree of the human HBV genotypes, which is remarkably consistent with their distribution patterns.

## Results

### Identity analyses in nucleotide and amino acid sequences

We selected 3 ancient HBV genomes from Germany for analyses here, 7074 year old Karsdorf from LBK culture in Lower-Saxony, 5353 year old Sorsum from Funnelbeaker culture from Lower-Saxony, and 4488 year old RISE563 from Bell Beaker culture in Osterhofen-Altenmarkt (Krause-Kyora et al. 2018; Muhlemann et al. 2018). The other Bronze age samples were not studied as one (RISE254) was very close to RISE563 and the others had many sequence gaps. In nucleotide identity, the three ancient samples were all closer to each other than to any present day samples, and the highest identity was between Sorsum and RISE563 (Table 1).

**Table 1.**
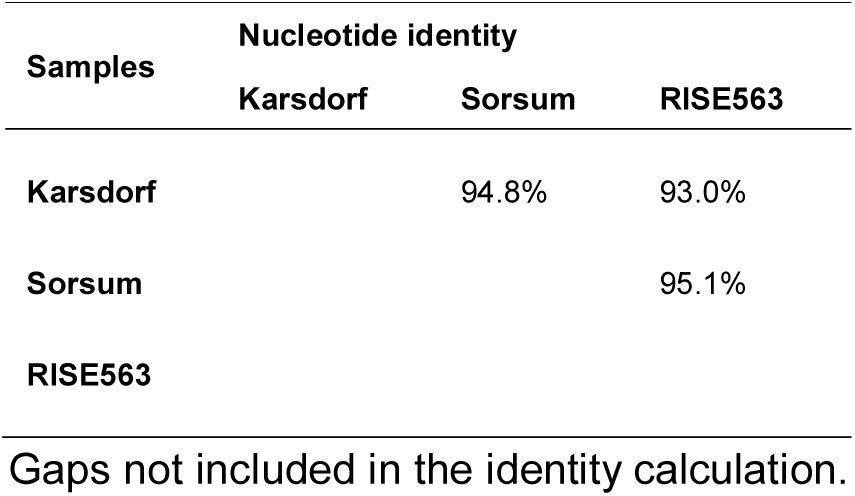
Nucleotide identity among the three ancient HBV genomes

We searched the Genbank protein database to identify the closest present day HBV genome to the ancient HBVs in amino acid identity in the polymerase, the largest open reading frame in the HBV genome (832-845 amino acids). Upon identifying the closest, we also examined its identity to the ancient HBV in other proteins, pre S, X, and core proteins (Table 2). Present day HBV isolates closest to the ancient HBVs in nucleotide identity were not found to be the closest in amino acid identity in the polymerase. For example, the Karsdorf sample was closest to a chimpanzee HBV (accession AB032433) in nucleotide sequence but a human HBV (HE981175, genotype G) in amino acid sequence in the polymerase. However, in other proteins, the closest to the ancient HBVs were all present day NHP HBVs (Table 2).

**Table 2.** Amino acid identities in the polymerase among ancient and present day HBV genomes.

| Proteins | Species | Karsdorf (7074) |  | Sorsum (5353) |  | RISE563 (4488) |  |
| --- | --- | --- | --- | --- | --- | --- | --- |
|  |  | Access. | Identity | Access. | Identity | Access. | Identity |
| Polymerase | Human | HE981175 | 92.9% | EU239218 | 92.5% | EU239218 | 90.8% |
|  | Chimp. | AB032433 | 92.6% | AB032433 | 92.5% | AB032433 | 90.9% |
|  | Gorilla | AJ131567 | 90.6% | AJ131567 | 90.6% | AJ131567 | 89.4% |
|  | Orangutan | AF193863 | 88.2% | AF193863 | 88.3% | AF193863 | 87.4% |
|  | Gibbon | AJ131571 | 91.3% | AJ131571 | 89.8% | AJ131571 | 88.7% |
|  | Karsdorf | Karsdorf |  | Karsdorf | 95.5% | Karsdorf | 93.5% |
|  | Sorsum | Sorsum | 95.5% | Sorsum |  | Sorsum | 94.8% |
|  | RISE563 | LT992443 | 93.5% | LT992443 | 94.8% | LT992443 |  |
| Pre S | Human | HE981175 | 91.1% | EU239218 | 92.3% | EU239218 | 91.7% |
|  | Ape | AB032433 | 92.6% | AB032433 | 95.6% | AJ131567 | 91.5% |
| X prot. | Human | HE981175 | 83.1% | EU239218 | 87.4% | EU239218 | 85.3% |
|  | Ape | AB032433 | 84.4% | AB032433 | 90.9% | AJ131567 | 91.7% |
| Core prot. | Human | HE981175 | 94.2% | EU239218 | 91.7% | EU239218 | 92.6% |
|  | Ape | AB032433 | 99.3% | AJ131567 | 96.1% | AJ131567 | 98.7% |

Note: Approximate age in years of ancient HBV samples is indicated next to sample name. The closest human HBV to the ancient HBV in polymerase identity was selected for comparison. Gaps were excluded in the identity calculation. The
underlined accession numbers indicate the closest modern HBV to the ancient HBV in nucleotide identity as previously published.

The 3 ancient HBV genomes were also closest to each other in polymerase amino acid identity than to any other present day samples (Table 2). However, different from the nucleotide result, the highest amino acid identity in polymerase was between Karsdorf and Sorsum. As Sorsum differs from RISE563 in both time periods and locations while only in time periods from Karsdorf, Sorsum is expected to be a closer relative of Karsdorf and hence to have more similarity in slow changing sites (amino acid) with Karsdorf. On the other hand, as Sorsum was 1721 years apart from Karsdorf, ~2 fold more than its time difference with RISE563 (865 years), Sorsum is expected to have more genetic distance from Karsdorf due to fast changing sites as may be reflected in the nucleotide sequence. Together, these results showed significant disconnect between amino acid and nucleotide sequences in revealing genetic relationships among HBVs.

While results in Table 2 showed clear affinity of Karsdorf with human HBV, Sorsum showed equal affinity with human and chimpanzee HBVs and RISE563 showed slightly more affinity to chimpanzee than to human HBV, indicating some uncertainty regarding the informative nature of the full length polymerase protein. The polymerase is composed of 4 domains, terminal protein, non-conserved spacer, reverse transcriptase, and RNase H. Upon examining the HBVdb database (https://hbvdb.ibcp.fr/) (Hayer et al. 2013), together with our own alignment analyses, we found that the amino acid region corresponding to the reverse transcriptase and RNase H domains are more conserved or slow evolving (343-844 aa for genotype G starting with VNL). We therefore tested this 501 aa region to see if it may show better results than the full length polymerase in linking ancient HBVs with human rather than NHP (Table 3). Again, all three ancient HBVs showed closer identity, but to a greater degree, to human HBVs than to NHP HBVs. While Karsdorf was again closest to Sorsum, it was closer to a present day human HBV (HE981175) than to the ancient HBV RISE563, indicating clear HBV genetic continuity from the time of Karsdorf to present time and the more informative nature of the 501 aa slow region of the polymerase.

**Table 3.** Amino acid identity among ancient and present day HBV genomes in the slow region (501 aa)

| Species | Karsdorf (7074) |  | Sorsum (5353) |  | RISE563 (4488) |  |
| --- | --- | --- | --- | --- | --- | --- |
|  | Access. | Identity | Access. | Identity | Access. | Identity |
| Human | HE981175 | 96.0% | HE981175 | 95.4% | EU239218 | 93.9% |
| Chimp. | <u>AB032433</u> | 93.8% | <u>AB032433</u> | 95.2% | AB032433 | 93.5% |
| Gorilla | AJ131567 | 93.8% | AJ131567 | 95.2% | <u>AJ131567</u> | 93.7% |
| Orangutan | AF193863 | 92.0% | AF193863 | 93.2% | AF193863 | 92.1% |
| Gibbon | AJ131571 | 93.2% | AJ131571 | 94.0% | AJ131571 | 92.2% |
| Karsdorf | Karsdorf |  | Karsdorf | 97.4% | Karsdorf | 94.6% |
| Sorsum | Sorsum | 97.4% | Sorsum |  | Sorsum | 96.2% |
| RISE563 | LT992443 | 94.6% | LT992443 | 96.2% | LT992443 |  |

As slow evolving DNAs are more likely to be in linear phase and hence more informative to phylogenetic relationships as explained by the MGD theory, we examined whether the 501 aa slow region of the polymerase is the slowest evolving among the protein genes in the HBV genome by comparing amino acid identity between human and orangutan HBV proteins (Supplementary Table S1). The 501 aa slow region of the polymerase was found to be the second most conserved, just slightly less conserved than the core protein. However, because the core protein was relatively short (178 aa), it is expected to be less informative than a longer protein with similar degree of conservation. Together with outgroup analyses (see below), we have found the 501 aa slow region of the polymerase to be the most informative to phylogenetic inferences of HBV strains.

### Outgroup inferences based on amino acid mutations

The above results raise the important question of which type of sequences may be most informative to HBV phylogeny. For viruses, different hosts may confer different physiological selection pressures which may result in viruses from different hosts to have drastic or non-conservative amino acid changes. Taking into account of non-conservative changes may thus be informative to phylogenetic relationships where an outgroup NHP HBV to two sister strains of human HBV is expected to show more non-conservative amino acid changes from the human HBVs.

To confirm the human rather than NHP affinity of the ancient HBV isolates as shown by the polymerase, we therefore performed protein alignment involving 3 strains, an ancient HBV, its closest human HBV, and its closest NHP HBV. It is expected that an outgroup should have a higher fraction of non-conservative or drastic amino acid changes among all mutations that led to differences between the outgroup and the other two sister strains. We examined those positions where the two sisters had the same residue while the outgroup was different. We tested each of the three compared HBV viruses as the candidate outgroup and obtained the fraction of drastic changes among all positions where the two sisters were the same while the outgroup was different (Table 4 and Supplementary Materials for details of this analysis). For the full length polymerase protein, RISE563 as the outgroup had a significantly smaller fraction of drastic changes than the gorilla HBV as the outgroup (0.33 vs 0.69, P = 0.004). This indicates that RISE563 and present day human HBV (EU239218) were sister strains while the gorilla HBV was the outgroup. Similar analyses showed that for Karsdorf and Sorsum samples, the NHP HBVs all showed the highest fraction of drastic changes (Table 4). We also performed the combined analysis where we first add up all the drastic changes of an outgroup (with the outgroup being ancient HBV, present day human HBV, or NHP HBV) and then calculated the fraction of drastic changes. The fraction of drastic changes in the NHP HBV when tested as the outgroup was significantly higher than either that in the ancient HBVs or the present day human HBVs when they were tested as the outgroup. We also obtained similar results for the slow region of the polymerase (Table 4). These results confirmed that ancient HBVs isolates grouped with human rather than NHP HBVs.

**Table 4.** Outgroup inferences from non-conservative (drastic) amino acid changes.

| Outgroup |  | Polymerase |  |  |  | Slow region (aa 343-844) |  |  |  |
| --- | --- | --- | --- | --- | --- | --- | --- | --- | --- |
|  |  | Mutations |  | Fraction | P value | Mutations |  | Fraction | P value |
| Isolates | Accession | Drastic | All | Drastic | Hu. v Ape | Drastic | All | drastic | Hu. v Ape |
| Karsdorf | NA | 5 | 16 | 0.31 | 0.02 | 5 | 10 | 0.50 | n.s. |
| Human | HE981175 | 9 | 24 | 0.38 | 0.02 | 4 | 9 | 0.44 | n.s. |
| Chimpanzee | AB032433 | 16 | 22 | 0.73 |  | 11 | 15 | 0.73 |  |
| Sorsum | NA | 7 | 16 | 0.44 | n.s. | 2 | 6 | 0.33 | n.s. |
| Human | EU239218 | 18 | 39 | 0.46 | n.s. | 5 | 18 | 0.28 | n.s. |
| Chimpanzee | AJ131575 | 26 | 39 | 0.67 |  | 4 | 7 | 0.57 |  |
| RISE563 | LT992443 | 8 | 24 | 0.33 | 0.004 | 2 | 6 | 0.44 | n.s. |
| Human | EU239218 | 16 | 32 | 0.50 | n.s. | 2 | 13 | 0.24 | n.s. |
| Gorilla | AJ131567 | 27 | 39 | 0.69 |  | 12 | 16 | 0.75 |  |
| Sum |  |  |  |  |  |  |  |  |  |
| Ancient human |  | 20 | 56 | 0.36 | <0.0001 | 9 | 22 | 0.41 | 0.03 |
| Present human |  | 43 | 95 | 0.45 | 0.0003 | 11 | 40 | 0.28 | 0.0002 |
| Ape |  | 69 | 100 | 0.69 |  | 27 | 38 | 0.71 |  |

In contrast, for the other three smaller size proteins of HBV genome, the Pre S protein, the X protein and the pre core protein, none was found informative in identifying an outgroup (Supplementary Table S2). When we did the same analysis by using these three proteins as concatenated single molecule, we also failed to identify any clear outgroup. The fractions of drastic changes in either ancient or present day human HBVs were similar to that of NHP and showed enrichment of non-conservative amino acid changes, which was unlike the case for the polymerase. Thus, the ancient HBVs did not group with either present day NHP or human in any of these proteins. That the observed changes were enriched for non-conservative amino acid mutations indicated functional adaptation or selection. Although ancient HBVs all showed slightly closer identities in these proteins to NHP HBVs, such weak affinity may be fortuitous.

### Phylogenetic relationships among HBV genotypes

The above results suggest that the slow region of the polymerase (aa 343-844) may be the most informative with regard to HBV phylogenetic relationships. We next used this region to reconstruct the phylogenetic tree for the 8 HBV genotypes by using the reference genomes for these genotypes (2 genomes for each genotype) as indexed by the HBVdb database. We first obtained the pairwise identities in the slow region among the 8 genotypes or 16 genomes and the average identity of each genotype to the other 7 genotypes (Table 5 and Supplementary Table S3). We also determined the lowest pairwise identity within each genotype by searching the Genbank database using the reference genomes and found D to have the lowest within genotype identity (461 aa), indicating that D has the largest within genotype genetic diversity among all genotypes. As D was also among the lowest in identity to all other genotypes (459.61, just slightly greater than the lowest H), D qualifies as the most basal genotype. We constructed a schematic diagram of the phylogenetic tree that best fits the data in Table 5 (Figure 1A). Relative to the existing tree (Figure 1B), the new tree is more consistent with the present day distribution pattern of the HBV genotypes (Figure 1C). In particular, consistent with expectations, the basal type D is the most widely distributed and most common in Northeast Asia and Russia/Siberia, and well positioned to split the first branch, genotype F and H, specific to the New World.

**Table 5.**
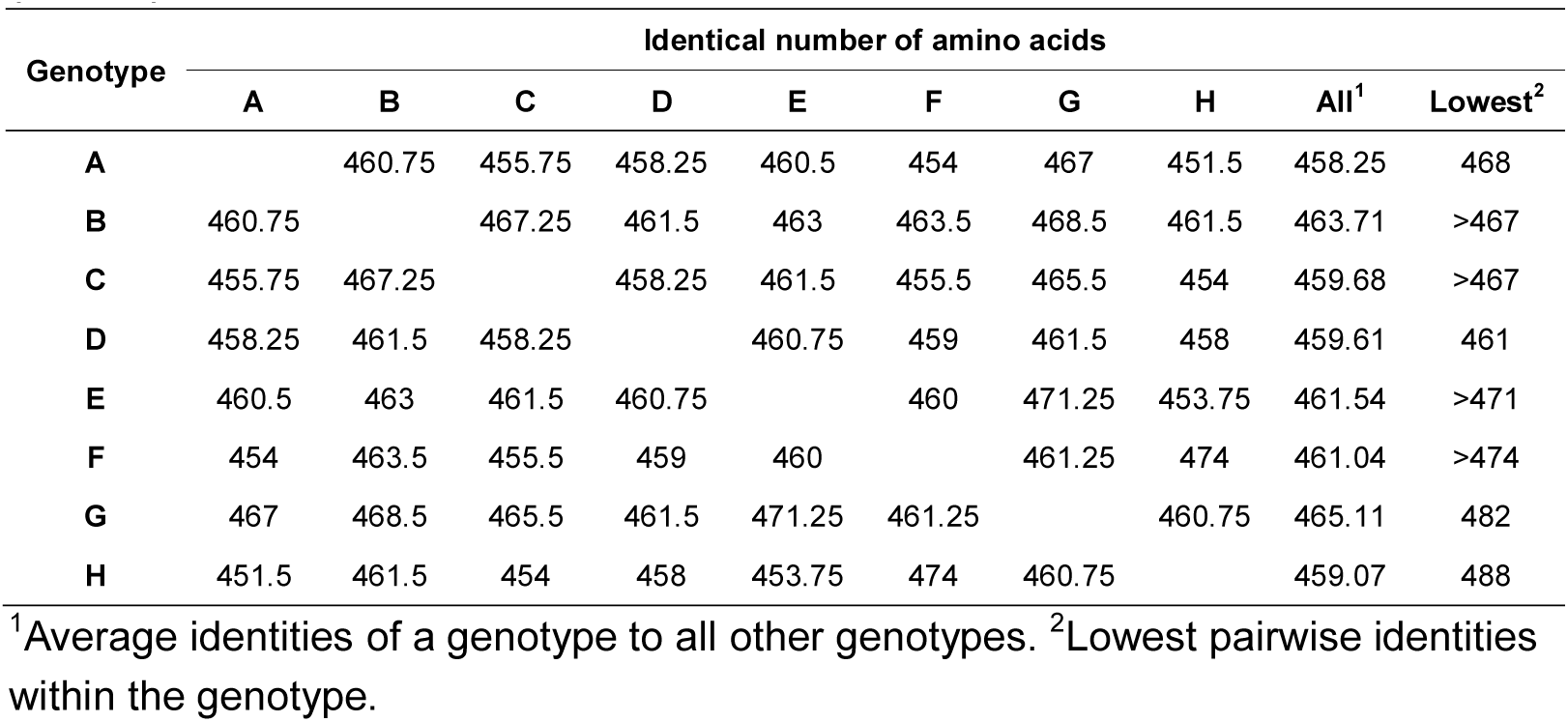
Average pairwise identities in the slow region of the HBV polymerase (501 aa)

**Figure 1.**
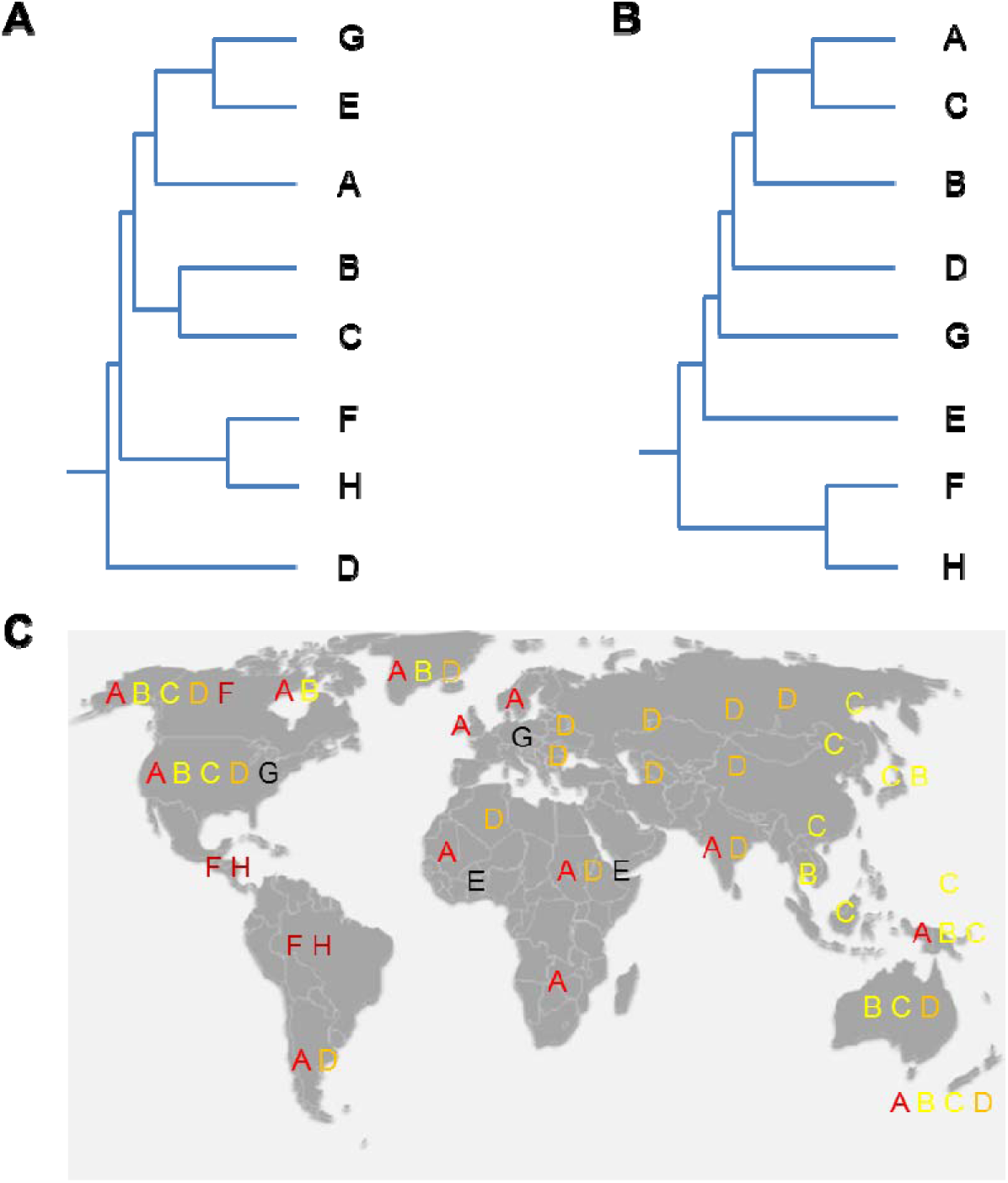
Phylogenetic tree of HBV genotypes. Branch lengths are relatively and roughly to scale. The tree is meant to be more of a schematic diagram. A. Tree built by using the slow region. B. Tree built by using nucleotide sequences as found in Muhlemann et al (Muhlemann et al. 2018). C. Distribution map of HBV genotypes.

Using the slow region of the polymerase, we also examined the relationships of selected NHP HBVs with the human reference genomes and found closer relative affinity of African NHP HBVs with genotype G and of Asian NHP HBVs with genotype C (Table 6). Although we only looked at one HBV genome each for each NHP species, the closest chimpanzee and gorilla in nucleotide sequence to the ancient human HBVs and an arbitrarily chosen orangutan and gibbon HBV, the relative affinity should hold for other NHP HBVs. We also examined the identity of ancient HBVs with the human HBV reference genomes and found all three to be most related to genotype G, with Sorsum and RISE563 relatively closer to genotype E than Karsdorf. The results suggest that Sorsum and RISE563 may be on their way diverging from genotype G to E. Based on the amino acid difference between Karsdorf and genotype G (501-479 = 22 amino acid) and the age of Karsdorf, we inferred the mutation rate in the slow region to be 2.0 x 10^-6^ aa per aa per year and the age of G at ~14500 years (14500 x 501 x 2.0 x 10^-6^ + 7500 x 501 x 2.0 x 10^-6^ = 22 aa). Given the distance between the basal genotype D and the other genotypes to be ~41 as well as the largest within genotype distance of 40 for D, we inferred an origination of human HBV at ~20500 years ago followed soon by the divergence of genotype F and H as people crossed the Bering Strait into the New World.

**Table 6.**
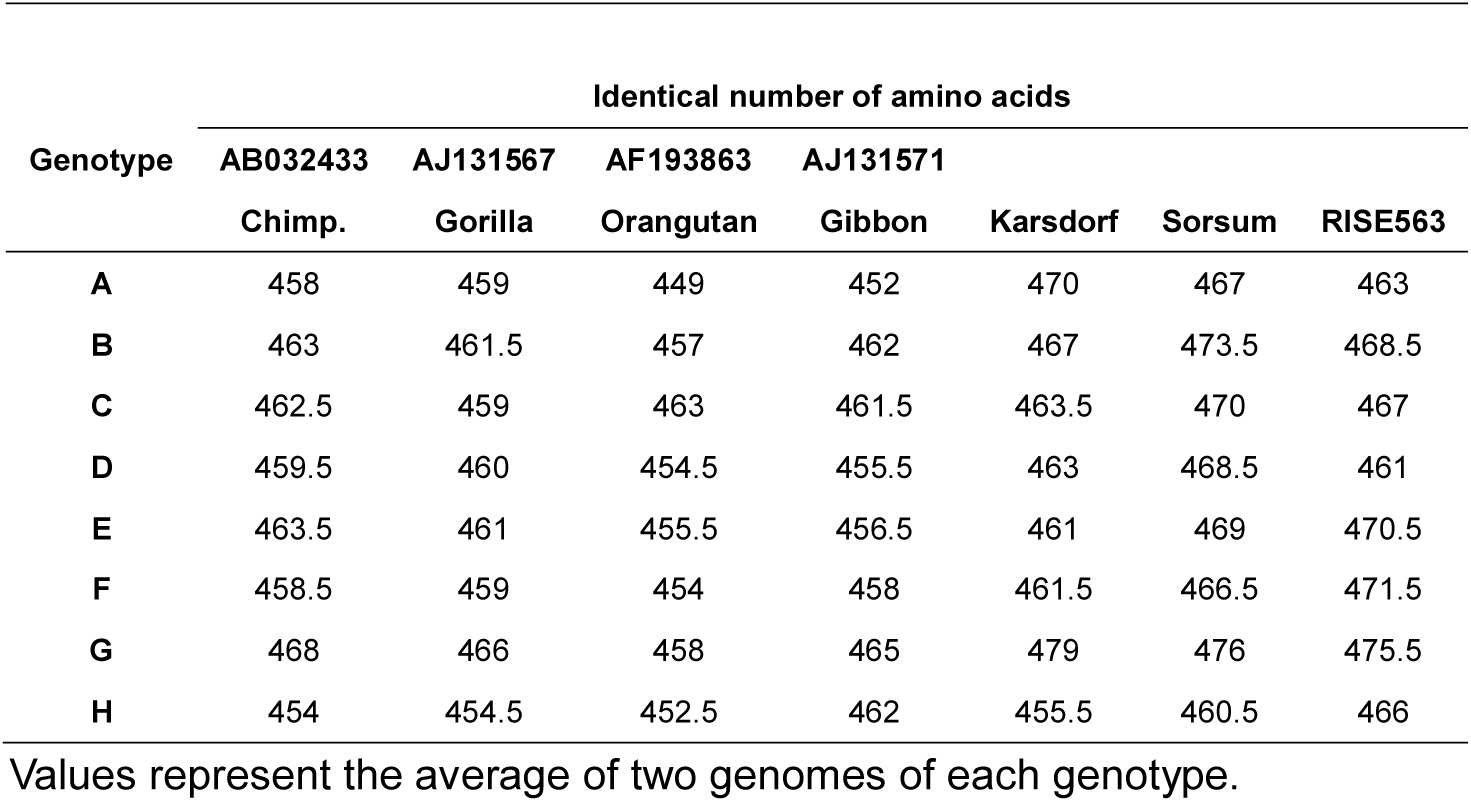
Identities between NHP and ancient HBV isolates with the reference genomes of human HBV genotypes

## Discussion

Our results suggest genetic continuity in the last seven Millennia for human HBV and establish a more informative method for building phylogenetic trees for HBVs. Our findings on the ancient HBV isolates here, rather than the conclusions from previous analyses (Krause-Kyora et al. 2018; Muhlemann et al. 2018), appear more consistent with a priori expectation on the ancient ancestors. That the oldest Karsdorf sample (7074 years ago) found in Germany was closest to present day genotype G that is also common in Germany, France and the United States is also consistent with expectation and further supports the conclusion of genetic continuity in HBVs in the last 7000 years (Sunbul 2014). That Sorsum and RISE563 samples were closely related to Karsdorf indicates also an affinity of these two isolates with genotype G although they were relatively more related to genotype E than Karsdorf. E is the most closely related to G among present HBV isolates. As E is found today only in Africa (Andernach, Hubschen, Muller 2009), our finding here is in line with the known migration of Europeans into Africa during the last 5000 year period (Fregel et al.2018).

The closer nucleotide distance of Sorsum with RISE563 than with Karsdorf may reflect linear phase distance whereas the closer amino acid distance of Sorsum with Karsdorf may reflect continuity in housekeeping functions. That Karsdorf was most similar to gorilla HBV in nucleotide sequence but to present day human HBV in amino acid sequence in the polymerase or the slow region may reflect continuation in the housekeeping physiology of the viruses and merely fortuitous similarities in nucleotide sequences reflecting some shared environmental adaptation. The closer similarity of ancient HBVs with NHP HBVs in non-polymerase proteins may reflect non-housekeeping adaptive roles in these proteins: the ancient HBV non-polymerase proteins in the ancient human hosts are expected to be more similar to NHP proteins given that the primitive life style of ancient humans may be less different from that of NHP.

The three ancient samples differ from genotype G however in that they lacked the 36 nucleotide insertion in the core gene region that is specific to HBV-G. This indicates that the insertion of this 36 bp sequence may have taken place later than 7000 years ago.

Genotype D is known to have the largest worldwide distribution, which is consistent with its basal position in the new tree here and its widest within genotype genetic diversity as found here. D shares the 33 nucleotide deletion in the pre S region with the NHP HBVs and the ancient HBVs, again indicating its more primitive nature. As all present day genotypes other than D lacks the 33 nucleotide deletion, we can infer that this deletion occurred only in the last few thousand years and independently in each genotype. As African NHP HBVs were closer to G while Asian NHP HBVs were closer to C, there was likely transmission of human HBVs into NHPs via migration of G from Europe into Africa and migration of C in the Asia mainland where it is most common today into the islands in Southeast Asia.

The new HBV phylogenetic tree appears more consistent with the reality genotype distribution around the globe than the existing ones and has implications for the origin of the virus. That the first split was between D and the rest followed by the split of F and H indicates that HBV may have originated in a region covering Northeast Asia and Siberia where D may have originated. It then went to either the West to give rise to G, E, and A or the South to give rise to B and C. The estimated age of the human HBV of ~20500 years corresponds to the period of Last Glacial Maximum (LGM), indicating that human population expansion post LGM may have played a role in the origin of HBV (Tallavaara et al. 2015).

The study here demonstrates the informative nature of the slow clock method in resolving the longstanding uncertainty on HBV origin and evolution. While it is widely known that different regions of a virus genome may produce different phylogenetic trees, it remains uncertain as to which region or whether the whole genome is the most informative to phylogenetic inferences. The MGD theory and the study here illustrate the informative nature of the slow evolving regions. The method should be generally applicable to evolutionary studies of other viruses.

## Materials and Methods

Present day HBV sequences were retrieved from Genbank and HBVdb database. Ancient HBV genomes were downloaded using previously published accession numbers. Alignments in nucleotide and amino acid sequences were done by blast.Fisher’s exact test was used to estimate p values, 2 tailed.

## Acknowledgements

We thank Ben Krause-Kyora and Julian Susat for sharing the sequences of ancient human HBV samples. This work is supported by the National Natural Science Foundation of China grant 81171880, the National Basic Research Program of China grant 2011CB51001, and the Furong Scholars program (S. H.).

## Author contributions

XL, YZ, SH designed the study and performed data analyses. SH wrote the first draft and all authors contributed to the final version.

## Supplementary Materials

### Original data on outgroup analyses for Karsdorf using the polymerase amino acid sequence

Transcriptase starts at 343 VNI

Karsdorf polymerase protein (had 33 nt deletion)

MPLSYQHFRRLLLLDDEAGPLEEELPRLADEGLNHRVAEDLNLQLPNVSIPWTHKVGNFT

GLYSSTVPVFNPXWQTPSFPDIHLHQDIINKCEQFVGPLTVNEKRRLQLVMPARFYPNST

KYLPLEKGIKPYYPDNVVNHYFQTRHYLHTLWKAGILYKRETTRSASFCGSPYSWEQELQ HGA

KXPFHKQSSRILSRPPVGPSVQSKYQQSRLGFQSQQGPLARGQQGRSWSIRARVHPTARR

PFGVEPSVSGHTNNIAS-------------------KRHSSSGHAVE

IPPNSARSQSEGPVFSCWWLQFRNSKPCSEYCLSHI

343 VNLLEDWGPCTEHGKHHIRIPRTPARVTGGVFLVDKNPHNTTESRLVVDFSQFSRG

STRVSWPKFAVPnlqsltnllssnlswl

sIDVSAAFYHIPLHPAAMPHLLVGSSGLSRYVARLSSDSRILDHQHGTMQNMHDSCSRNL

FVSLMLLYKTFGRKLHLYSHPIILGFRKIPMGVGLSPFLLAQFTSAICSVVRRAFPHCLA

FSYMDDVVLGAKTVQHLESLYTAVTNFLLSLGIHLNPNKTKRWGYSLNFMGYVIGSWGTL

PQDHIIQKIKQCFRKLPVNRPIDWKVCQRIVGLLGFAAPFTQCGYPALIVIPLYACIQAKQA

FTFSPTYKAFLCKQYLNLYPVARQRPGLCQVFADATPTGWGLVIVIGHQRIVIRGTFVAPLPIH

TAELLAACFARSRSGANLIGTDNSVVLSRKYTSFPWLLGCAANWILRGTSFVYVPSALNP

ADDPSRGHLGLCRPLLRLSYQPTTGRTSLYAVSPSVPSHLPDRVHFASPLHVTWK

Karsdorf nucleotide sequences:

NNNNNNNACATTCCACCAAACTCTGCAAGATCCCAGAGTGAGGGGCCTGTATTTTCCTGC

TGGTGGCTCCAGTTCAGGAACAGTAAACCCTGTTCCGAATACTGCCTCTCACACATCGTC

AATCTTCTCGAGGACTGGGGACCCTGCACCGAACATGGAAAACATCACATCAGGATTCCT

AGGACCCCTGCTCGCGTTACAGGCGGGGTTTTTCTTGTTGACAAAAATCCTCACAATACC

ACAGAGTCTAGACTCGTGGTGGACTTCTCTCAATTTTCTAGGGGGAGCACCCGTGTGTCT

TGGCCAAAATTCGCAGTCCCCAACCTCCAATCACTCACCAACCTCCTGTCCTCCAACCTG

TCCTGGCTATCGCTGGATGTGTCTGCGGCGTTTTATCATATTCCTCTTCATCCTGCTGCT

ATGCCTCATCTTCTTGTTGGTTCTTCTGGACTATCAAGGTATGTTGCCCGTCTGTCCTCT

GATTCCAGGATCCTCGACCACCAGCACGGGACCATGCAGAACATGCACGACTCCTGCTCA

AGGAACCTCTTTGTATCCCTCATGTTGCTGTACAAAACCTTCGGACGGAAATTGCACCTG

TATTCCCATCCCATCATCCTGGGCTTTCGGAAAATTCCTATGGGAGTGGGCCTCAGTCCG

TTTCTCCTGGCTCAGTTTACTAGCGCCATTTGTTCAGTGGTTCGCAGGGCTTTCCCCCAC

TGTTTGGCTTTCAGTTATATGGATGATGTGGTTTTGGGGGCCAAGACTGTACAACATCTT

GAGTCCCTTTACACCGCTGTTACTAATTTTCTTTTGTCTTTGGGCATACATTTAAATCCC

AACAAAACAAAAAGATGGGGTTATTCCCTAAACTTCATGGGTTATGTAATTGGAAGTTGG

GGAACATT GCCACAGGAT CACATT ATT CAGAAAAT CAAACAAT GTTT CAGAAAACT CCCT

GTTAACAGACCTATTGATTGGAAAGTATGTCAAAGAATTGTGGGTCTTTTGGGCTTTGCC

GCCCCTTTTACACAATGTGGTTATCCAGCATTAATGCCTTTATATGCATGTATACAAGCT

AAGCAGGCTTTCACTTTCTCGCCAACTTACAAGGCCTTTCTGTGTAAACAATATTTGAAC

CTTTACCCCGTTGCCCGGCAACGGCCAGGTCTGTGCCAAGTGTTTGCTGATGCAACCCCC

ACTGGCTGGGGCTTGGTCATGGGCCATCAGCGCATGCGTGGAACCTTTGTGGCTCCTCTG

CCGATCCATACTGCGGAACTCCTAGCCGCTTGTTTTGCTCGCAGCAGGTCTGGAGCAAAC

CTTATTGGGACTGATAATTCTGTTGTCCTCTCCCGGAAATATACATCATTTCCATGGCTG

CTAGGCTGTGCTGCCAACTGGATCCTGCGCGGGACGTCCTTTGTTTACGTCCCGTCAGCG

CTGAATCCTGCGGACGACCCCTCTCGGGGCCACTTGGGGCTTTGCCGCCCCCTTCTCCGT

CTGTCGTACCAGCCGACCACGGGGCGCACCTCTCTTTACGCGGTCTCCCCGTCTGTGCCT

TCTCATCTGCCGGACCGTGTGCACTTCGCTTCACCTCTGCATGTTACATGGAAACCGCCG

TGAACGCCCCCCGGAACCTGCCAAGGGACTTACATAAGAGGACTCTTGGACTCTCAGCAATGTCAACAACCAAGATT

GAGACATACTTCAAAGACTGTGTATTTGAGGAGTGGGAGGAATCAGGCAAGGACACCAGGTTAATGACCTTTGTATTAGGAGGCTGTAGGCATAAATTGGTCT

GTTCACCAGCACCATGTAACTTTTTCACCTCTGCCTAATCATCTCTTGTTCATGTCCTAC

TGTTCAAGCCTCCAAGTTGTGCCTTGGGTGGCTTTTGGGCATGGACATTGACCCATATAA

AGAATTTGGAGCTACTGTTGAGTTGCTCTCCTTTTTGCCTTCTGACTTTTTTCCTTCGGT

CCGCGATCTTCTCGACACCGCCTCAGCTCTGTACCGGGAAGCCTTAGAGTCTCCAGAGCA

TTGTT CACCAAATCACACAGCACTCAGGCAAGCTGTTCTGT GTTGGGGTGAGTT AATGAC

CTTGGCTTCCTGGGTGGGCAACAATTTGGAAGATCNNNNNNNNNNNNNNNNNNNNNNNNN

NNNNNNNNNNNNNNNNNNATGGGTTTAAAAATAAGGCAACTATTGTGGTTTCACATTTCC

TGTCTTACTTTTGGAAGAGAAACGGTCCTTGAGTATTTGGTGTCTTTTGGAGTGTGGATT

CGCACTCCTCCCGCTTACAGACCACCAAATGCCCCTATCTTATCAACACTTCCGGAGACT

ACTGTTGTTAGACGACGAGGCAGGTCCCCTAGAAGAAGAACTCCCTCGCCTCGCAGACGA

AGGTCTCAATCACCGCGTCGCAGAAGATCTCAATCTCCAGCTTCCCAATGTTAGTATTCC

TTGGACTCACAAGGTGGGAAACTTTACGGGGCTTTATTCTTCTACTGTTCCTGTCTTTAA

CCCTNNCTGGCAAACTCCTTCTTTTCCTGATATTCATTTGCATCAAGATATCATTAACAA

ATGCGAACAATTTGTGGGCCCTCTTACAGTAAATGAAAAACGAAGATTACAGTTAGTTAT

GCCTGCTAGATTTTACCCTAACTCTACAAAATATTTGCCCCTAGAGAAAGGCATAAAGCC

TTATT ATC CAGATAAT GTGGTT AAT CATT ACTTCCAAACCAGACATT ATTT ACAT ACTCT

ATGGAAGGCGGGCATCTTATATAAAAGAGAGACAACACGTAGCGCCTCATTTTGTGGGTC

ACCATATT CTTGGGAACAAGAGCTACAGCATGGGGCAGAATCTNNNNNNNNNNNNNNNNN

NNNNNNNNNNNNNNNNNNNNNNNNNNNNNNNNNTGGAGAAANAACCTTTCCACAAGCAAT

CCTCTAGGATTCTTTCCCGACCACCAGTTGGACCCAGCGTTCAGAGCAAATACCAACAAT

CCAGATTGGGATTTCAATCCCAACAAGGACCCTTGGCCAGAGGCCAACAAGGTAGGAGCT

GGAGCATTCGGGCCAGGGTTCACCCCACCGCACGGAGGCCTTTTGGGGTGGAGCCCTCAG

TCTCAGGGCATACTAACAACATTGCCAGCAAGAGACACTCATCCTCAGGCCATGCAGTGG AA

Karsdorf aligned with Human CCK86662 and chimp AB032433

Region: 306-832 aa

For the 501 amino acid (343-844) slow region, the number of drastic changes among total changes are the following:

Karsdorf 5/10 (among 10 changes that differ between Karsdorf and human/chimp, 5 are drastic)

Human 4/9 (among 9 changes that differ between Human and Karsdorf/chimp, 4 are drastic)

Chimp 11/15 (among 15 changes that differ between Chimp and Karsdorf/human, 11 are drastic)

#### Conclusion

Chimp is the outlier to the clade containing Karsdorf and human HBV CCK86662.

### Alignments performed by blastp

Kars. 10

IPPNSARSQSEGPVFSCWWLQFRNSKPCSEYCLSHIVNLLEDWGPCTEHGKHHIRIP

RTP 189

IPP+S +SQS+GPVFSCWWLQFR+S+PCS+YCLSH+VNLL+DWGPCTEHG+HHIRIPRTP

Human 306

IPPSSTKSQSQGPVFSCWWLQFRDSEPCSDYCLSHLVNLLQDWGPCTEHGEHHIRIPRTP 365

chimp 306

KGPVFSCWWLQFRNIEPCSEYCLSHL<u>VSLLDDWGPCNEHGEHHIRIPRTP</u>

Informative changes are highlighted in yellow. Green positions show all three to have different residues, which are not informative and not counted.

1. Karsdorf as candidate outgroup
2. Human as candidate outgroup
3. Chimp as candidate outgroup

1. 0/3 (among 3 changes that differ between Karsdorf and human/chimp, 0 are drastic)
2. 0/2 (among 2 changes that differ between Human and Karsdorf/chimp, 0 are drastic)
3. 2/2 (among 2 changes that differ between Chimp and Karsdorf/human, 2 are drastic)

343 slow region

1. 0/1
2. 0/0
3. 2/2

Kars. 190 ARVTGGVFLVDKNPHNTTESRLVVDFSQFSRGSTRVSWPKFAVPnlqsltnllssnlswl 369

ARVTGGVFLVDKNPHNT ESRLVVDFSQFSRGS

RVSWPKFAVPNLQSLTNLLSSNLSWL

Sbjct 366

ARVTGGVFLVDKNPHNTAESRLVVDFSQFSRGSARVSWPKFAVPNLQSLTNLLSSNLSWL 425

Sbjct 366

ARITGGVFLVDKNPHNTAESRLVVDFSQFSRGSTRVPWPKFAVPNLQSLTNLLSSNLSWL 425

1. 1/1
2. 1/1
3. 1/1

Kars. 370

sIDVSAAFYHIPLHPAAMPHLLVGSSGLSRYVARLSSDSRILDHQHGTMQNMHDSCSRNL 549

SLDVSAAFYHIPLHPAAMPHLLVGSSGLSRYVARLSSDSRILDHQHGT+QN+HDSCSRL

Sbjct 426 SLDVSAAFYHIPLHPAAMPHLLVGSSGLSRYVARLSSDSRILDHQHGTLQNLHDSCSRQL

Sbjct 426

SLDVSAAFYHLPLHPAAMPHLLVGSSGLSRYVARLSSNSRILDHQHGTMQNLHDSCSRNL 485

1. 0/1
2. 1/2
3. 0/1

Kars. 550 FVSLMLLYKTFGRKLHLYSHPIILGFRKIPMGVGLSPFLLAQFTSAICSWRRAFPHCLA 729

+VSLMLLYKTFGRKLHLYSHPIILGFRKIPMGVGLSPFLLAQFTSAICSWRRAFPHCLA Sbjct 486 YVSLMLLYKTFGRKLHLYSHPIILGFRKIPMGVGLSPFLLAQFTSAICSWRRAFPHCLA 545

Sbjct 486 FDSLMLLYKTFGRKLHLYSHPIIMGFRKIPMGVGLSPFLLAQFTSAICSVVRRAFPHCLA 545

1. 0/0
2. 0/1
3. 1/2

Kars. 730

FSYMDDVVLGAKTVQHLESLYTAVTNFLLSLGIHLNPNKTKRWGYSLNFMGYVIGSWGTL 909

FSYMDDVVLGAK+VQHLESLYTAVTNFLLSLGIHLNPNKTKRWGYSLNFMGYVIGSWGTL

Sbjct 546

FSYMDDVVLGAKSVQHLESLYTAVTNFLLSLGIHLNPNKTKRWGYSLNFMGYVIGSWGTL 605

Sbjct 546

FSYMDDVVLGAKSVQHLESLYTAVTNFLLSLGIHLNPNKTKRWGYSLHFMGYVIGSWGTL 605

1. 0/1
2. 0/0
3. 1/1

Kars. 910 PQDHIIQKIKQCFRKLPVNRPIDWKVCQRIVGLLGFAAPFTQCGYPALMPLYACIQAKQA 1089

PQ+HI QKIKQCFRKLPVNRPIDWKVCQRI GLLGFAAPFTQCGYPALMPLYACIQAKQA

Sbjct 606 PQEHITQKIKQCFRKLPVNRPIDWKVCQRITGLLGFAAPFTQCGYPALIVIPLYACIQAKQA 665

Sbjct 606 PQEHIVQKIKNCFRKLPVNRPIDWKVCQRIVGLLGFAAPFTQCGYPALMPLYACIQAKQA 665

1. 0/1
2. 0/0
3. 0/1

Kars. 1090

FTFSPTYKAFLCKQYLNLYPVARQRPGLCQVFADATPTGWGLVMGHQRMRGTFVAPLPIH 1269

FTFSPTYKAFLCKQY+NLYPVARQRPGLCQVFADATPTGWGL +GHQRMRGTFVAPLPIH

Sbjct 666

FTFSPTYKAFLCKQYMNLYPVARQRPGLCQVFADATPTGWGLAIGHQRMRGTFVAPLPIH 725

Sbjct 666

FTFSPTYKAFLSQQYSTLYPVARQRSGLCQVFADATPTGWGLVMGHQRMRGTFVAPLPIH 725

1. 1/2
2. 1/2
3. 3/3

Kars. 1270 TAELLAACFARSRSGANLIGTDNSVVLSRKYTSFPWLLGCAANWILRGTSFVYVPSALNPTAELLAACFARSRSGA LIGTDNSVVLSRKYTSFPWLLGCAANWILRGTSFVYVPSALNP 1449

Sbjct 726 TAELLAACFARSRSGAKLIGTDNSVVLSRKYTSFPWLLGCAANWILRGTSFVYVPSALNP

785

Sbjct 726 TAELLAACFARSRSGAKLIGTDNSVVLSRKYTSFPWLLGCAANWILRGTSFVYVPSALNP

785

1. 1/1
2. 0/0
3. 0/0

Kars. 1450 ADDPSRGHLGLCRPLLRLSYQPTTGRTSLYAVSPSVPSHLPDRVHFASPLHVTWK

1614

ADDPSRG LGLCRPLLRL + PTTGRTSLYAVSPSVPSHLPDRVHFASPLHVTWK

Sbjct 786 ADDPSRGRLGLCRPLLRLPFLPTTGRTSLYAVSPSVPSHLPDRVHFASPLHVTWK 840

Sbjct 786 ADDPSRGRLGLYRPLIRLLFQPTTGRTSLYAVSPSVPSHLPVRVHFASPLHVAWR 830

1. 1/2
2. 1/1
3. 3/4

